# Adult-born hippocampal neurons undergo extended development and are morphologically distinct from neonatally-born neurons

**DOI:** 10.1101/702746

**Authors:** John Darby Cole, Delane Espinueva, Désirée R. Seib, Alyssa M. Ash, Matthew B. Cooke, Shaina P. Cahill, Timothy O’Leary, Sharon S. Kwan, Jason S. Snyder

## Abstract

During immature stages, adult-born neurons pass through critical periods for survival and plasticity. It is generally assumed that by 2 months of age adult-born neurons are mature and equivalent to the broader neuronal population, raising questions of how they might contribute to hippocampal function in old age when neurogenesis has declined. However, few have examined adult-born neurons beyond the critical period, or directly compared them to neurons born in infancy. Here, we used a retrovirus to visualize functionally-relevant morphological features of 2- to 24-week-old adult-born neurons in male rats. From 2-7 weeks neurons grew and attained a relatively mature phenotype. However, several features of 7-week-old neurons suggested a later wave of growth: these neurons had larger nuclei, thicker dendrites and more dendritic filopodia than all other groups. Indeed, between 7-24 weeks, adult-born neurons gained additional dendritic branches, grew a 2^nd^ primary dendrite, acquired more mushroom spines and had enlarged mossy fiber presynaptic terminals. Compared to neonatally-born neurons, old adult-born neurons had greater spine density, larger presynaptic terminals, and more putative efferent filopodial contacts onto inhibitory neurons. By integrating rates of cell birth and growth across the lifespan, we estimate that adult neurogenesis ultimately produces half of the cells and the majority of spines in the dentate gyrus. Critically, protracted development contributes to the plasticity of the hippocampus through to the end of life, even after cell production declines. Persistent differences from neonatally-born neurons may additionally endow adult-born neurons with unique functions even after they have matured.

**SIGNIFICANCE STATEMENT:** Neurogenesis occurs in the hippocampus throughout adult life and contributes to memory and emotion. It is generally assumed that new neurons have the greatest impact on behavior when they are immature and plastic. However, since neurogenesis declines dramatically with age, it is unclear how they might contribute to behavior later in life when cell proliferation has slowed. Here we find that newborn neurons mature over many months in rats, and end up with distinct morphological features compared to neurons born in infancy. Using a mathematical model, we estimate that a large fraction of neurons is added in adulthood. Moreover, their extended growth produces a reserve of plasticity that persists even after neurogenesis has declined to low rates.

## INTRODUCTION

Morphological and physiological studies of adult-born dentate gyrus (DG) neurons suggest that adult neurogenesis may play an important role in hippocampal function. During the first 2-8 weeks of neuronal development, adult-born neurons are less constrained by GABAergic inhibition and display greater afferent and efferent synaptic potentiation (Snyder et al., 2001; Schmidt-Hieber et al., 2004; Ge et al., 2007; Gu et al., 2012; Marín-Burgin et al., 2012; Chancey et al., 2013). They have greater excitability, which enables them to be recruited despite immature innervation by cortical inputs (Mongiat et al., 2009; Dieni et al., 2013). During defined windows of time they also undergo experience-dependent survival and innervation by excitatory and inhibitory neurons, and recruit GABAergic interneurons at higher rates (Epp et al., 2007; Tashiro et al., 2007; Anderson et al., 2010; Bergami et al., 2015; Vivar et al., 2015; Alvarez et al., 2016). The transient nature of these unique properties suggests that adult-born neurons have the greatest impact on circuits and behavior when they are in an immature critical period (Aimone et al., 2009; Kim et al., 2012; Snyder and Cameron, 2012).

While adult-born neurons eventually acquire features of developmentally-born neurons (Laplagne et al., 2006; Stone et al., 2011), the extent to which they are similar is unclear because few studies have examined adult-born neurons beyond the traditional critical window of ∼2-6 weeks. There is evidence that even old adult-born neurons have a unique capacity for experience-induced morphological growth and immediate-early gene expression (Lemaire et al., 2012; Tronel et al., 2015). Additionally, studies that have characterized adult-born neurons at older ages typically have not directly compared them to neurons born in development, making it difficult to conclude whether adult-born neurons are fundamentally similar or distinct from developmentally-born granule neurons. Work that has examined neurons born at different stages of life has found differences in the rate of maturation (Overstreet-Wadiche et al., 2006; Trinchero et al., 2017), neuronal survival (Dayer et al., 2003; Cahill et al., 2017; Ciric et al., 2019), immediate early gene expression (Imura et al., 2018; Ohline et al., 2018), morphology and physiology (Kerloch et al., 2018; Save et al., 2018). Thus, there appears to be an ontogenetic basis for cellular heterogeneity in the DG (Snyder, 2019).

In most mammals, neurogenesis declines approximately 90% between young and mid-adulthood and, by old age, newborn neurons are scarce (Lazic, 2012). If new neurons are particularly important during a brief window of immaturity, can neurogenesis make a significant contribution to hippocampal function later in life, when few neurons are added? This question is important because the DG is highly-vulnerable to age-related pathology (Geinisman et al., 1986; Small et al., 2004; Yassa et al., 2010) and the extent of neurogenesis in human aging is unclear (Charvet and Finlay, 2018; Kempermann et al., 2018; Paredes et al., 2018; Snyder, 2019; Flor-García et al., 2020). One possibility is that adult-born (or later-born) neurons may continue to mature and display developmental plasticity beyond the traditional critical period. The fact that 4-month-old adult-born neurons display enhanced spatial learning-induced morphological plasticity (Lemaire et al., 2012) suggests that old adult-born neurons may still have “room to grow” later in life when fewer neurons are being generated. Second, protracted neurogenesis may contribute to the functional heterogeneity of the DG by producing distinct types of neurons at different stages of life (Snyder, 2019). In this way, neurons born in adulthood may mature to become distinct from neurons born in development, and may therefore offer unique functions even in old age, when neurogenesis rates have declined. To test these possibilities we used a tdTomato-expressing retrovirus to visualize the detailed morphology of various-aged DG neurons in rats. By examining dendrites, spines and presynaptic terminals, we find that adult-born neurons continue to develop, and become morphologically-distinct from neonatally-born neurons, over an extended period of 24 weeks.

## METHODS

### Animals

Male Long Evans rats were used in the following experiments. They were bred and housed in the animal facility of the Department of Psychology using wild-type breeders from Charles River Canada. All manipulations were conducted according to guidelines of the Canadian Council on Animal Care and with protocols approved by the UBC Animal Care Committee. Rats were weaned at 21 days of age, pair housed (cages 48 x 27 x 20 cm) in male-only colony rooms that were separate from breeders, and given ad libitum access to water and rat chow. Cages were kept in on a 12-hour light/dark cycle, with the light cycle starting at 9:00 am. All manipulations were conducted in the light phase.

### General experimental design

An overview of the experimental design is provided in Fig. 1. The general approach was to use a tdTomato-expressing retrovirus to birthdate cohorts of DG neurons that were either born in neonates (P1) or adults, and enable visualization and quantification of their morphological properties. To birthdate animals, pregnant dams were checked daily for litters and P1 was defined as the first day that pups were observed in the cage. Most groups were examined when rats were 16 weeks old; retrovirus was injected at different times prior to this endpoint to examine neurons at different stages of cellular development, and to compare them to neurons born in the neonatal period. Neuronal ages were 2w (16 rats), 4w (8 rats), 7w (13 rats) and 16w (15 rats; neonatal-born). An additional cohort of adult-born neurons was allowed to survive until 24w (7 rats), and this was the only group in the main experiment that was examined at a different animal age (32w). In a follow-up experiment we injected groups of rats at 8 weeks of age (n=4) or 14 weeks of age (n=5) and examined cells 7 weeks later.

**Figure 1:**
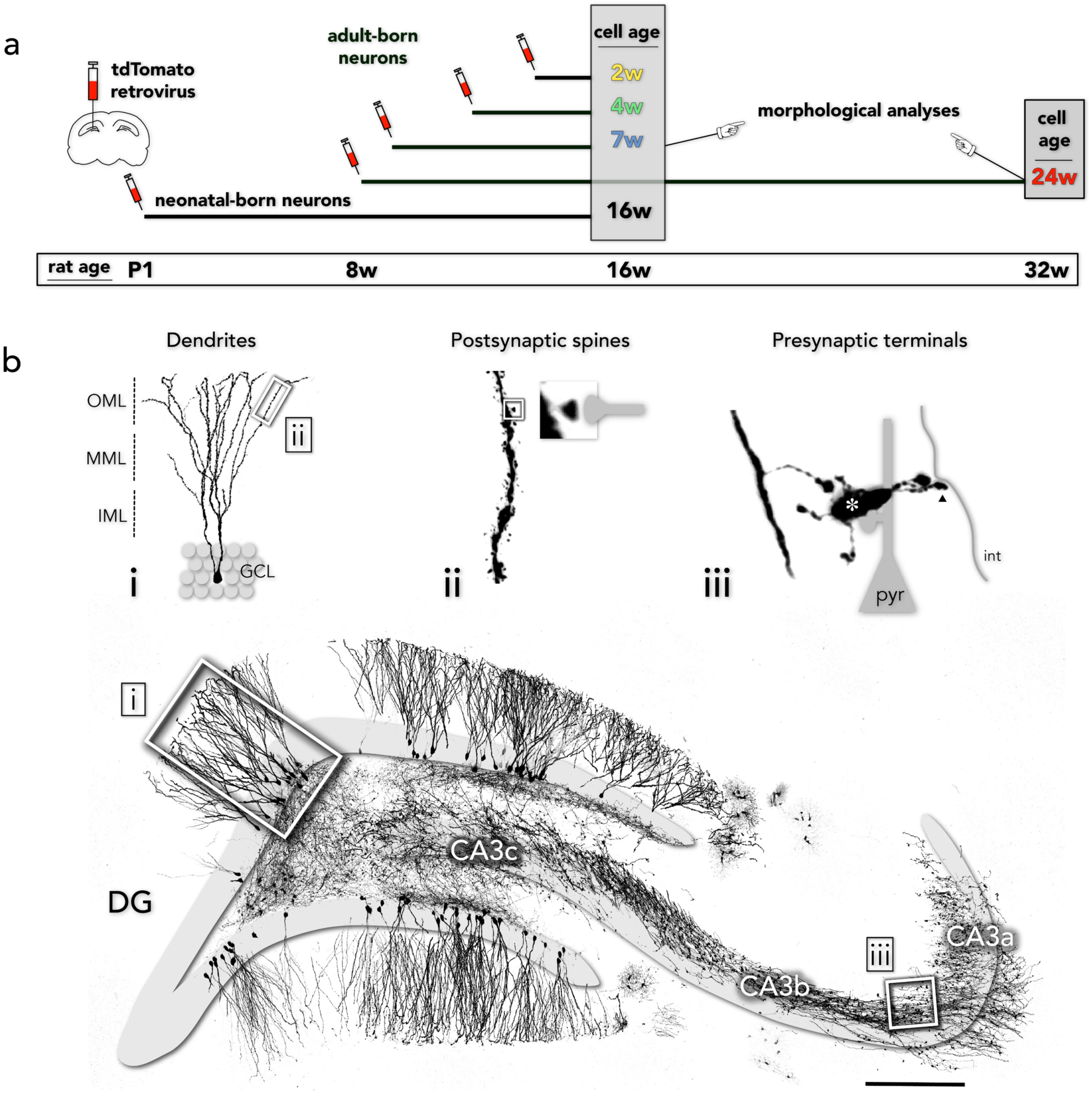
Experimental design. a) Timeline: tdTomato-expressing retrovirus was injected into the DG of rats to birthdate granule neurons and visualize their morphological features. All groups were examined when rats were 16 weeks of age, except one cohort of adult-born cells that were allowed to mature for 24 weeks (rat age 32 weeks) and a follow-up experiment that examined 7-week-old neurons born in 8- and 14-week-old rats (not shown here). b) The large low magnification confocal image shows retrovirally-labelled, 24-week-old neurons in the dorsal hippocampus; granule and CA3 pyramidal cell layers were traced from DAPI^+^ principal cell nuclei. Insets highlight the morphological features of DG neurons that were investigated: i) dendritic trees; ii) spines; iii) presynaptic mossy fiber boutons (*), which target CA3 pyramidal neurons (pyr), and filopodial terminals (arrowhead), which target inhibitory interneurons (int). OML, outer molecular layer; MML, middle molecular layer; IML, inner molecular layer; GCL, granule cell layer. Scale bar, 500 μm.

Some animals in the 2w, 4w, 7w and 16w-neonatal groups were additionally trained for 1 day in a spatial water maze, 1 week prior to endpoint, to examine possible experience-dependent effects on structural morphology. However, since effects of water maze training were minimal, cells from control and trained rats were pooled for most analyses.

### Retrovirus production

The retroviral vector used in this study was derived from a Moloney Murine Leukemia-Virus (MMLV), in which tdTomato expression is driven by a Ubiquitin (Ubi) promoter as described previously (van Praag et al., 2002). Retroviral Ubi-tdTomato (MMLV-tdTomato; kindly provided by Dr. Shaoyu Ge) and VSV-G (kindly provided by Dr. Ana Martin-Villalba) plasmids were transfected in HEK293-GP cells (kindly provided by Dr. Diane Lagace) using polyethylenimine. Retrovirus was harvested 2 and 3 days after transfection, followed by ultracentrifugation (2 h at 89,527g). Viral titers ranged from 0.8 to 30 x 10^4^ colony forming units/ml.

### Stereotaxic retrovirus injection into the dorsal dentate gyrus

MMLV-tdTomato was injected into the DG of rats according to sterile surgical procedures approved by the UBC animal care committee. Rats were anaesthetized with isoflurane, given Ketoprofen (5 mg/kg), local bupivacaine and lactated ringers (10 ml/kg) every hour during the surgery. For adult surgeries, heads were levelled and fixed in a stereotaxic frame (Kopf, Tujunga, CA) and retrovirus injections were made at - 4.0 mm posterior, ±3.0 mm mediolateral and −3.5 mm ventral relative to bregma. One μl of retrovirus was injected into each hemisphere, at a speed of 200 nl/min, using a 30 gauge Hamilton needle and a microsyringe pump (World Precision Instruments, Sarasota, FL). The needle remained in place for 5 min after the injection to allow the retrovirus to diffuse. For neonatal surgeries, pups were anesthetized with isoflurane, manually secured in the stereotaxic apparatus, and injected with 500 nl of retrovirus into the dorsal hippocampus (position estimated by eye, relative to lambda) over ∼30 sec (Mathon et al., 2015).

### Spatial water maze training

Experimental and cage control animals from the 2w, 4w, 7w and 16w-neonatal groups were habituated to handling for 1 week prior to the start of training. Trained rats were subjected to 8 trials in a standard spatial water maze. The pool diameter was 2 m, water was 20°C and made opaque with white nontoxic tempera paint, and the platform (10 cm in diameter) was submerged 1 cm below the surface of the water. Distal 2D and 3D cues were located 1-3 meters away, on the room walls, providing the spatial information needed to learn to navigate to the hidden platform.

### Tissue processing

Animals were deeply anesthetized with isoflurane and transcardially perfused with 120 ml of 4% paraformaldehyde (PFA) in phosphate-buffered saline (PBS; pH 7.4). Extracted brains were immersed in 4% PFA for an additional 48 hours at 4°C and were then transferred to PBS. Using a vibratome, brain sections were cut at 100 μm and kept in their rostro-caudal sequence in 48-well plates to facilitate reconstruction of neurons across multiple sections. Slices were then incubated in 0.1 M citric acid in a 99°C water bath for 15 minutes, were then washed in PBS, and then incubated in 10% PBS-TX with 3% horse serum (ThermoFisher Scientific, cat# 16050122) for 30 minutes. Sections were then incubated in rabbit anti-RFP (1:1000; Rockland cat# 600401379) with 10% PBS-TX with 3% horse serum, for 72 hours, on a shaker, at 4^°^C. Sections were then washed with PBS-TX, incubated with donkey anti-rabbit Alexa-Fluor 555 (1:250; ThermoFisher Scientific, cat# A31572) for 60 minutes at room temperature. After another PBS wash, slices stained with DAPI (1:1000 in PBS) for 5 minutes. Slices were washed with PBS four more times before being serially mounted onto slides (Fisherbrand Superfrost Plus) and coverslipped with PVA-DABCO.

### Imaging and morphological analyses

For all morphological analyses, images of tdTomato^+^ neurons in the dorsal DG (−2.8 mm to −4.8 mm relative to Bregma) were acquired with a Leica SP8 confocal microscope. Dendrite and spine analyses were restricted to the suprapyramidal blade, to avoid variation due to blade differences (Claiborne et al., 1990). All animals with successful bilateral or unilateral labelling were analyzed. While there was a range of viral titers, lower and higher titer virus was distributed across groups and all animals within a given group had comparable tdTomato staining intensity. tdTomato immunostaining was sufficiently robust in all groups to enable reliable visualization and quantification, though staining intensity varied from cell to cell, and tended to be weaker in immature cells. To maintain approximate consistency across images, laser power and gain were adjusted to maintain a comparable dynamic range and ensure that brightness of the faintest spines was *∼*25-30, which was just visible by eye (8-bit images, ranging 0-255). To achieve this, the laser power ranged from *∼*0.2 to 3% and the gain ranged from *∼*15 to 60%. Unless stated otherwise, analyses and measurements were performed on the z-stacks to accurately distinguish fine morphological details from each other and from background noise that can interfere with signals (e.g. particularly in maximum intensity projections).

For dendritic analyses, images 1024 x 1024 pixels in size and at a z-resolution of 1.25 μm were acquired with a 25x, water-immersion lens (NA 0.95) at 1x zoom. Granule cells were imaged across adjacent sections to obtain the full dendritic tree. Neuronal dendrites were traced in Image J with the Simple Neurite Tracer plugin (Longair et al., 2011). The full dendritic tree (i.e. across multiple sections) was included for analyses of total dendritic length and dendritic branching order (1°, 2°, 3° etc., using the Neuroanatomy plugins for ImageJ). Sholl analyses of dendritic branching were performed on individual sections that contained ≥ 70% of the total dendritic length of a given neuron (see details in Results). Dendrite thickness was measured from spine/protrusion images (see below) and calculated as the average of 3 thickness measurements taken at both ends and the middle of the 30-70 μm segment. A single segment was measured from each of the inner, middle and outer molecular layers per cell.

Dendritic protrusion images were acquired with a glycerol-immersion 63x objective, at 1024 x 1024 pixels, 0.75 μm z-resolution, and at 5x zoom. Segments 30 μm to 70 μm in length, from cells in the suprapyramidal blade, were sampled from the inner, middle and outer molecular layers (molecular layer divided into 3 zones of equal width, approximating the terminal zones for hilar, medial entorhinal and lateral entorhinal axons, respectively; Fig. 1b). Typically, the same neurons were sampled in all 3 layers and, for total protrusion analyses, values were averaged. Protrusions, obvious elongations that extend approximately perpendicular from the dendrite, were counted with the ImageJ Cell Counter plugin. They were categorized according to morphological classes that vary with maturity (Toni et al., 2007; Berry and Nedivi, 2017): filopodia (immature, thin extensions that lack a bulbous head and are typically devoid of synapses), thin spines (putative post-synaptic spines that have a bulbous head and a thin neck), stubby spines (short and lacking a spine neck; spines with this appearance were included in protrusion density calculations but were not separately analyzed) and mushroom spines (mature, stable spines with synapses; here defined as those with a large head of ≥ 0.6 μm in diameter).

Large mossy fiber boutons (MFBs) were imaged with a glycerol-immersion 63x objective, at 1024 x 1024 pixels, 1 μm z-resolution, and at 5x zoom. MFBs were sampled randomly within CA3a, CA3b, and CA3c and were identified by their large, irregular shape and (typically) associated filopodial extensions (Claiborne et al., 1986; Acsády et al., 1998). Cross-sectional area was measured on maximum projections of stacked images using Image J. Filopodia, protrusions from the bouton between 1 μm and 25 μm in length (Restivo et al., 2015), were analyzed from z-stacks.

Soma and nuclear sizes were measured from the neurons that were imaged for dendritic tree analyses, from the z-plane that had the largest tdTomato^+^ cell body and associated DAPI^+^ nucleus, respectively (typically the middle plane of the cell).

### Modelling neurogenic plasticity across the lifespan

We developed a mathematical model to estimate the cumulative effects of neurogenesis throughout the lifespan, focusing on dendritic length and spine numbers. Using MATLAB, we identified functions that effectively fit age-related changes in neurogenesis and the growth of dendrites and spines as adult-born neurons matured. Peak DG neurogenesis was defined as 166,000 cells born on postnatal day 6 (Schlessinger et al., 1975; Cahill et al., 2017) and rates of adult neurogenesis were estimated by scaling published ^3^H-Thy^+^ and BrdU^+^ cell counts (Altman and Das, 1965; Kuhn et al., 1996) relative to this peak, as we have done previously (Snyder, 2019). Neurogenesis rates were estimated by fitting a double Gaussian function to cell count data from P30 onwards; only adult data (P56 onwards) are included in the model. We did not correct for inflated counts due to redivision of labelled precursor cells since: 1) embryonic and perinatal datasets were limited to heavily-labelled granule neurons, thereby largely excluding cells that would be labelled due to redivision, 2) inflation due to redivision in adulthood will be approximately offset by death of immature neurons (redivision causes ∼2x increase in labelled cells (Cameron et al., 1993), death removes ∼1/2 of cells in rats (Snyder et al., 2009)). The total (bilateral) granule cell population was fixed at 2.4 million cells (West et al., 1991), and 1 existing neuron (born before P56) was removed for each adult-born neuron that was added (Dayer et al., 2003; Cahill et al., 2017). For adult-born neurons, age-related increases in dendritic length and mushroom spines were fit with power functions and total spines (thin+mushroom) were fit with a sigmoidal function. We then integrated dendrite and spine growth functions for all neurons born between P56 and P730 to predict morphological consequences of neurogenesis in adulthood (code: https://github.com/MatthewBCooke/NeurogenesisFunctions). Dendrite lengths and spine densities for neurons born prior to adulthood were fixed at levels observed in P1-born neurons.

### Experimental Design and Statistical Analyses

Neuronal morphology varied substantially, even within the same animal. To retain these details, and compare subpopulations of neurons of the same age, we performed most analyses at the level of the structure of interest (i.e. cell, bouton), except where indicated otherwise. In some instances we also compared animal averaged data (e.g. to ensure that analyses were not skewed by over representation of cells from an “outlier” animal). Morphological differences between different-aged neurons were typically assessed by standard or repeated measures ANOVA with Holm-Sidak post-hoc comparisons. Samples that were not normally distributed were log transformed prior to statistical analyses and, if distributions remained non-normal, the untransformed data were analyzed by a non-parametric Kruskal-Wallis test with post-hoc comparisons by Dunn’s test. All graphs show non-transformed data. Cells born in 8w-old vs 14w-old animals were compared by two-tailed, unpaired t-tests except for branch order patterns, which were compared by repeated measures ANOVA. Statistical analyses can be found in the main text for data that are not presented in the figures. For data that are presented in the figures, statistical analyses can be found in the figure legends. The underlying data for all analyses are provided as Figure 2-1. In all cases, significance was set at α = 0.05.

**Figure 2:**
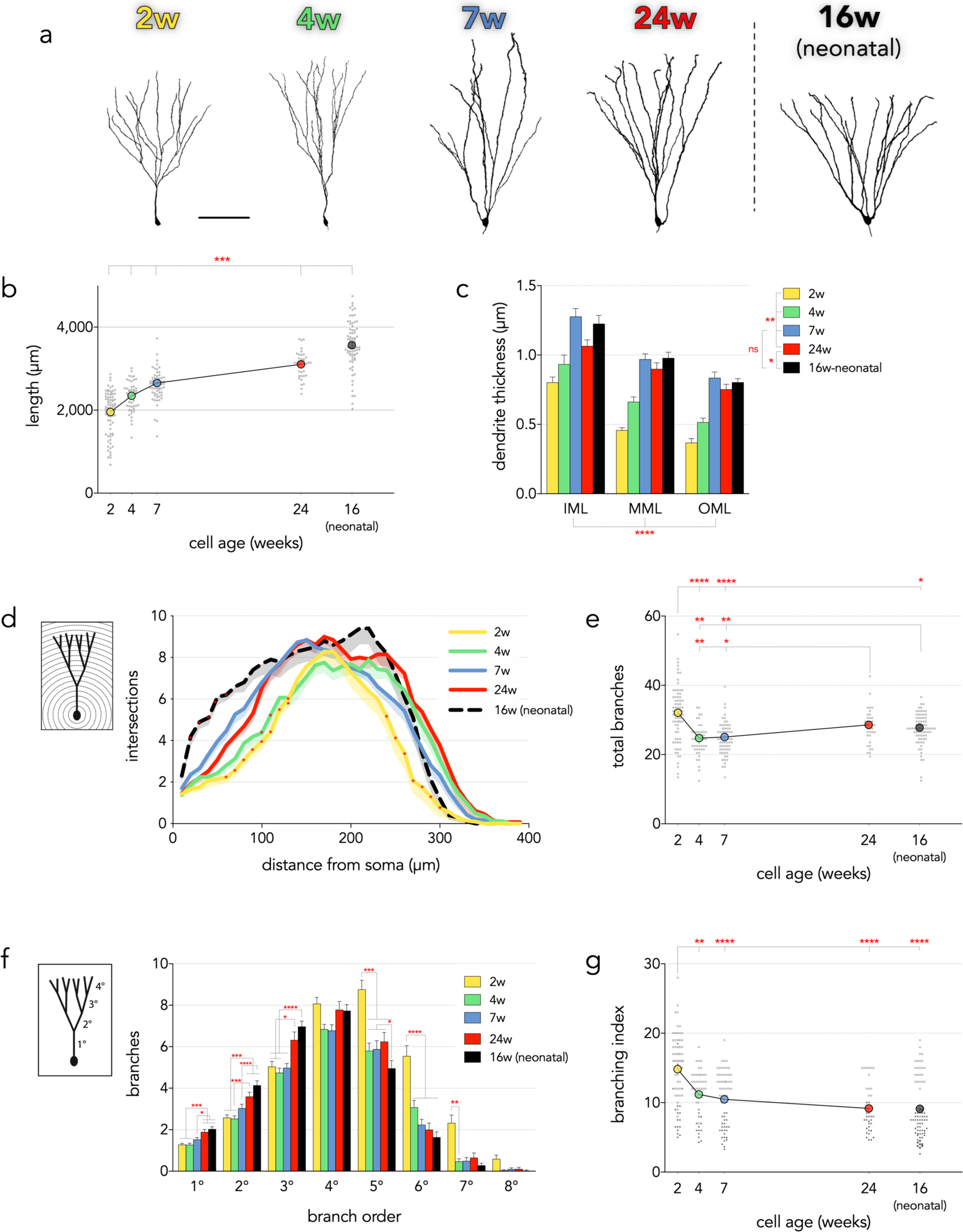
Dendritic structure of neonatally-born neurons and developing adult-born neurons. a) Representative confocal images of fully reconstructed dendritic trees. Scale bar, 100 μm. Examples of trifurcating dendrites in Movie 1. Raw data is provided as Figure 2-1. b) The total dendritic length of adult-born neurons increased from 2 to 24 weeks and remained shorter than neonatally-born neurons (F_4,296_=120, P<0.0001; ***P<0.001 for all group comparisons). Colored symbols indicate group means, small grey circles indicate dendritic lengths of individual neurons. c) Dendrites became thicker with increasing adult-born neuron age, with the exception that 7w cells had thicker dendrites than all adult-born groups, including 24w cells. Neonatally-born cells had thicker dendrites than 24w adult-born neurons. Dendrite thickness varied such that OML < MML < IML (repeated measures, mixed effects model; effect of cell age F_4,260_=52, P<0.0001; effect of molecular layer region, F_1.7, 402_=142, P<0.0001; cell x layer interaction, F_8,477_=0.9, P=0.5). d) Sholl analyses revealed distinct patterns of dendritic complexity across cell ages (effect of cell age F_4,85_=16, P<0.0001; cell age x dendritic subregion interaction F_116,2465_=2.9, P<0.0001; 10-300 μm analyzed since cells did not reliably extend beyond 300 μm in all groups). The total number of intersections was different between all groups (all comparisons P<0.05). Two-week-old neurons had fewer intersections than older adult-born neurons in both proximal and distal dendritic regions (60-130 μm, *P<0.05 vs 7w and 24w cells; 240-300 μm, *P<0.05 vs 24w cells). Neonatal-born neurons had more intersections in proximal dendritic regions (20-60 μm, *P<0.05 vs all other groups). Four-week-old neurons had fewer intersections than 7 and 24-week-old neurons at 100, 110 and 130 μm (*P<0.05). Lines connect mean values (not shown), shading indicates s.e.m. e) Two-week-old adult-born neurons had more dendritic branches than all other groups except 24w cells. After initial pruning from 2w to 4w, the number of branches increased from 7w to 24w (Kruskal-Wallis test, P<0.0001 followed by Dunn’s post-test). Symbols as in (b). f) Dendritic branching varied as a function of branch order and cell age (effect of cell age F_4,1682_=7.1, P<0.0001; effect of dendrite order F_7,1682_=394.1, P<0.0001, interaction F_28,1682_=9.0, P<0.0001). Neonatal-born neurons, and 24w-old adult-born neurons, had more lower-order branches than 2- 7w-old adult-born neurons. In contrast, young adult-born neurons (particularly 2w) had more high-order branches. Bars indicate mean ± s.e.m. g) The branching index (branch tips / # primary dendrites) was greater in 2w cells than all other cell ages (Kruskal Wallis test P<0.0001). White circles, grey squares, black triangles and grey circles indicate cells with 1, 2, 3 and 4 primary dendrites, respectively. The branching index was also greater in 2w cells than in other adult-born cells when only single primary dendrite cells were analyzed (Kruskal Wallis test P<0.0001; 2w vs 4w and 7w: P<0.0001, 2w vs 24w: P=0.02, 2w vs 16w-neonatal: P=0.4). *P<0.05, **P<0.01, ***P<0.001, ****P<0.0001.

## RESULTS

### Water Maze Behavior

Average latency to escape from the water maze decreased from 50 sec on trial 1 to 25 sec on trial 8, and there were no differences between groups (effect of trial, F_7,322_=8.8, P<0.0001; effect of cell age group F_3,46_=1.5, P=0.22). Average path length taken to escape also decreased across trials (1631 cm on trial 1 to 709 cm on trial 8) and was not different between groups (effect of trial, F_7,322_=11.7, P<0.0001; effect of cell age group F_3,46_=1.8, P=0.16).

### Minor effects of water maze training on neuronal morphology

Spatial water maze training over multiple days induces morphological and electrophysiological plasticity in adult-born neurons (Ambrogini et al., 2010; Tronel et al., 2010; Lemaire et al., 2012). Since the hippocampus is essential for remembering brief experiences (Feldman et al., 2010) and adult-born DG neurons show rapid changes in spine morphology following electrical stimulation (Ohkawa et al., 2012; Jungenitz et al., 2018), we hypothesized that a single session of water maze training may be sufficient to induce morphological plasticity in DG neurons. Contrary to our predictions, water maze training had minimal impact on the morphology of neonatally-born or adult-born neurons. These findings therefore do not contribute to the main conclusions of our study and, for our main analyses, data from trained and untrained rats are pooled. Nonetheless, we report the data from trained and untrained rats here:

Total dendritic length did not differ between control and water maze-trained rats (effect of training F_1,257_=1.7, P=0.19; training x cell age interaction F_3,257_=1.5, P=0.22).

Protrusion densities were greater in the inner molecular layer of water maze-trained rats but there was no difference between cell age groups (effect of training F_1,237_=5.0, P=0.03; training x cell age interaction F_3,237_=0.7, P=0.6). There was no effect of training on protrusion densities in the middle molecular layer or outer molecular layer (training effects P>0.25, interactions P>0.08). In the inner molecular layer, mushroom spine densities were greater in 7-week-old cells in water maze-trained rats (effect of training F_1,237_=2.8, P=0.1; training x cell age interaction F_3,237_=5.2, P=0.002; 7w cells in trained vs untrained rats: P=0.002). Water maze training increased mushroom spine densities in the middle molecular layer, but this was not different between cell age groups (effect of training F_1,235_=5.9, P=0.02, interaction F_3,235_=1.1 P=0.3). Water maze training did not significantly impact mushroom spine densities in the outer molecular layer (training and interaction effects both P>0.6).

Water maze training did not alter MFB size in CA3a or CA3b (training and interaction effects all P>0.07). In CA3c, MFBs were smaller in 2-week-old cells in water maze-trained rats (cell age x training interaction: F_3,195_=2.9, P=0.03; MFBs in trained vs untrained rats: P=0.006). Water maze training did not alter the number of filopodia/MFB, or the length of filopodia, in any CA3 subregion (training and interaction effects all P>0.11).

### Dendrites

Consistent with previous reports, the dendritic tree of adult-born neurons matured over several weeks (Fig. 2). Two and 4-week-old neurons had noticeably thinner and more irregular-shaped dendrites that often did not extend to the hippocampal fissure. By 7 weeks, dendrites were thicker, longer and, at even older ages, tips of dendrites often curved sideways upon approaching the hippocampal fissure. While less common than in younger cells, early-terminating dendrites (Fig. 2a) and thin, spine-poor dendritic segments (Fig. 4a) were also observed on 7- and 24- week-old neurons, suggesting the presence of immature processes and continued growth.

**Figure 3:**
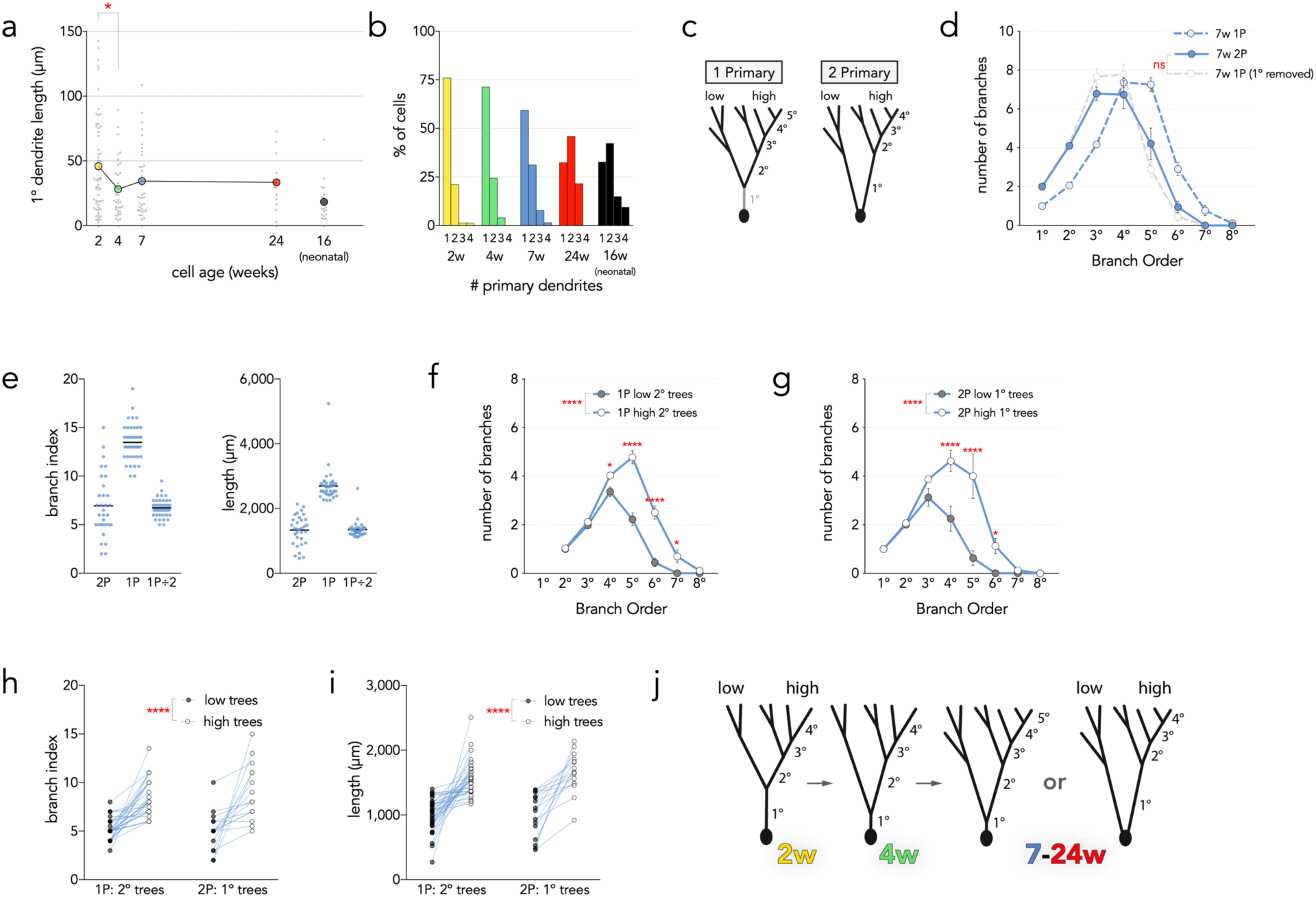
Unzipping model of primary dendrite formation. a) The primary dendrite shortens between 2-4 weeks of cell age (only includes cells with a single primary dendrite; F_4,167_=5.1, P=0.0007). b) The number of primary dendrites per cell, by age. c) Schematic of approach for comparing branch-tree morphology in cells with 1 vs 2 primary dendrites (1P vs 2P). By “removing” the primary dendrite from 1P cells their dendritic trees could be directly compared with 2P cells. “Low” trees had fewer branches and shorter total length than “high” trees (see below). d) Dendritic branch orders are similar in 7w-1P and 7w-2P cells, once the number of primary dendrites is accounted for (7w-2P vs 7w-1P with 1° dendrite removed, effect of cell type: F_1,35_=0.03, P=0.9; cell type x branch order interaction, F_6,210_=2.2, P=0.04, post hoc comparisons at each order all P>0.1). e) Primary dendritic trees on 7w-2P cells displayed more variable degrees of complexity than primary dendritic trees of 7w-1P cells, as measured by branch index and total length. f) 7w-1P cells had one 2º dendritic tree that branched significantly less (“low”) than the other (“high”; effect of tree type: F_1,35_=53, P<0.0001; tree type x branch order interaction: F_6,210_=24, P<0.0001). g) 7w cells with 2 primary dendrites had one 1º dendritic tree that branched significantly less than the other (effect of tree type: F_1,15_=16, P<0.01; tree type x branch order interaction: F_7,105_=9.8, P<0.0001). h) 7w cells had 2 main dendritic trees that differed in amount of branching, regardless of whether the cells had 1 or 2 primary dendrites (effect of tree type: F_1,50_=67, P<0.0001; tree type x cell type interaction: F_1,50_=1.0, P=0.3). i) “High” trees had greater total dendritic length than “low” trees, in both cells with 1 and 2 primary dendrites (effect of tree type: F_1,50_=71, P<0.0001; tree type x cell type interaction: F_1,50_=0.4, P=0.5). j) Unzipping model: The dendritic tree of adult-born neurons is inherently variable, with some “sub-trees” having more branches than others. Most adult-born neurons begin with a single primary dendrite, which shortens as the first branch point moves closer to the soma. In many cells the first branch point reaches the soma causing the transition from 1 to 2 primary dendrites, and differential complexity carries over as 2º-based subtrees become 1º-based subtrees.

**Figure 4:**
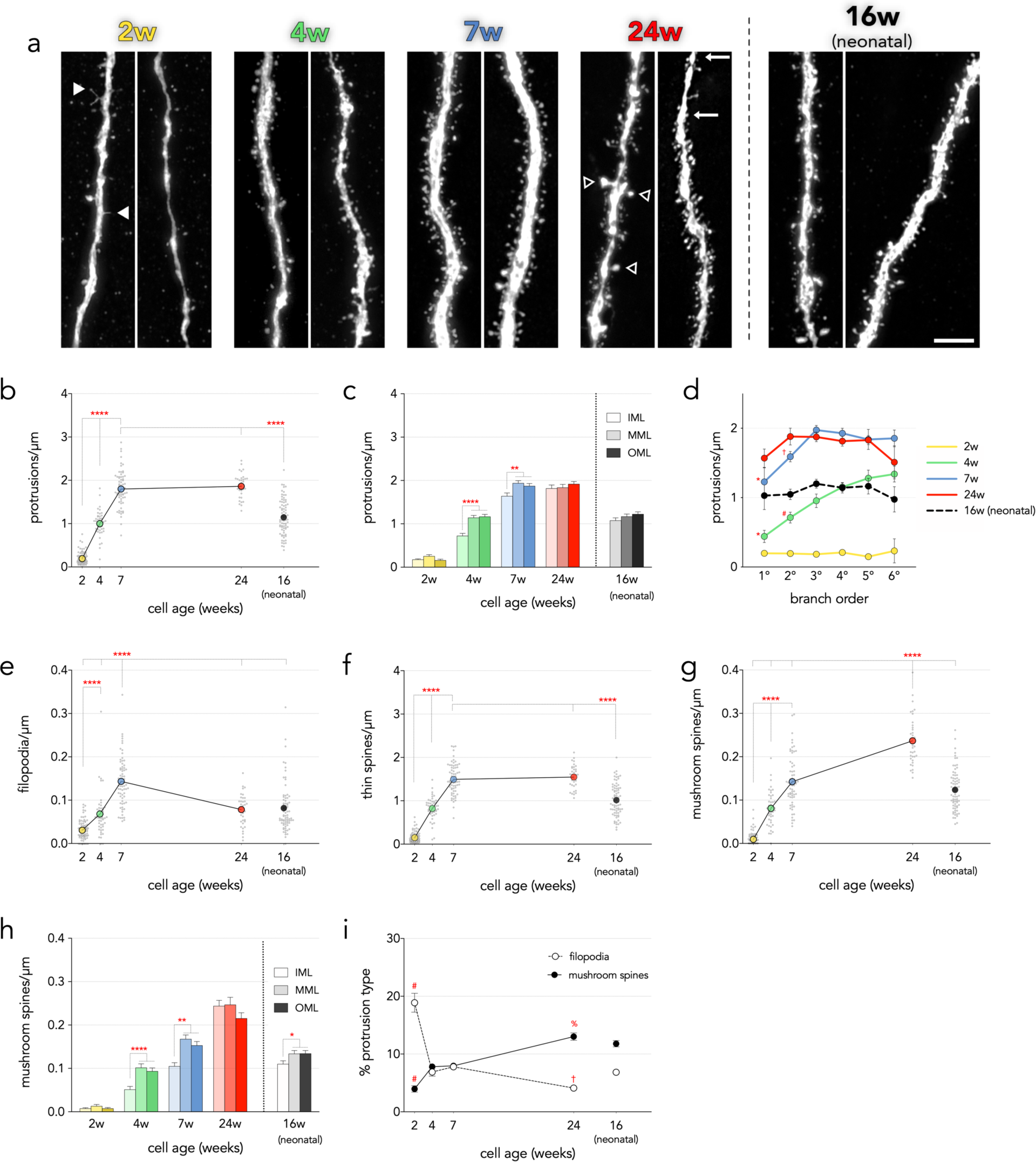
Spine densities in adult-born neurons reach and surpass that of neonatal-born neurons. a) Confocal images of spines/protrusions. Filled arrowheads in the 2 week example indicate filopodia; open arrowheads in the 24w example indicate mushroom spines; segment identified by arrows at 24w demonstrates region of low spine density in the distal tip of a dendrite in the outer molecular layer. Scale bar, 10 μm. b) Total protrusion densities increase with age in adult-born neurons, and plateau by 7 weeks at levels that are greater than neonatal-born neurons (F_4,314_=299, P<0.0001). c) Protrusion densities increased with cell age at a slower rate in the inner molecular layer than in the middle and outer molecular layers (cell age x layer interaction F_8,883_=3.5, P=0.0005). d) In immature neurons, protrusion densities were reduced in lower-order dendrites (branch order x cell age interaction F_20,615_=2.9, P<0.0001; 1° vs 3°,4°,5°,6° *P<0.05; 2° vs 4°,5°,6° ^#^P<0.01; 2° vs 3°,4° ^†^P<0.01). e) Filopodia densities increased from 2w to peak levels at 7w, and declined by 24w, which was not different that neonatally-born neurons (F_4,314=65_, P<0.0001). f) Thin spines made up the majority of protrusions and increased over 7 weeks to levels that were greater than neonatally-born neurons (F_4,314_=335, P<0.0001). g) Mushroom spine densities increased as adult-born neurons aged and, by 24 weeks, were greater than all other groups (F_4,314_=201, P<0.0001). h) Mushroom spine densities increased with adult-born neuron age at a slower rate in the inner molecular layer. Mushroom spine densities were also lower in the inner molecular layer of neonatally-born neurons (cell age x subregion interaction (F_8,883_=4.7, P<0.001). The proportion of filopodial protrusions was greatest in 2w cells and the proportion of mushroom spines was greatest in 24w cells (Kruskal-Wallis tests for both protrusion types, P<0.0001). Symbols and bars indicate means, error bars indicate s.e.m. IML: inner molecular layer, MML: middle molecular layer, OML: outer molecular layer. *P<0.05, **P<0.01, ***P<0.001, ****P<0.0001, ^#^P<0.001 vs same protrusion type at all other ages, ^%^P<0.0001 vs same protrusion type at 4w and 7w, ^†^P<0.05 vs same protrusion type at 4w, 7w, 16w-neonatal.

To determine the timeframe of growth of adult-born neuron dendrites, dendritic lengths were measured in their entirety, across sections. The total number of cells examined were: 76 (2w), 49 (4w), 65 (7w), 37 (24w), 74 (16w-neonatal). Among adult-born cells, dendritic length increased from 1954 μm at 2 weeks to 3105 μm at 24 weeks (Fig. 2b). There was greater dendritic growth at younger cell ages; from 2-4 weeks there was a net increase of 392 μm (28 μm/day) whereas over the much longer interval of 4-24 weeks there was only an additional 759 μm of growth (4 μm/day). Neonatally-born neurons had an average total dendritic length of 3565 μm; this was greater than all adult-born neuron groups, though populations overlapped.

Group means that were calculated from animal averaged data differed by less than 1% from means that were calculated by pooling all cells of a given age category. The same pattern of group differences were also observed when groups were comprised of animal averaged data rather than individual cell data (F_4,55_=57, all cell ages significantly different from one another: 0.05 > P < 0.0001).

As an additional index of dendritic growth, we measured the width of dendrites in the inner, middle and outer molecular layers (Fig. 2c). Dendrite thickness doubled between 2-7 weeks (0.5 to 1.0 μm) and then decreased slightly by 24w, which was thinner than 16w-neonatal neurons. Consistent with theoretical predictions that dendrites taper to optimize current transfer (Bird and Cuntz, 2016), dendritic thickness decreased from the inner to middle to outer molecular layers, and this did not vary across cell age.

Dendritic branching patterns change as adult-born neurons develop (Kerloch et al., 2018) and their precise morphology likely determines the strength and integration of synaptic inputs from different pathways (Spruston, 2008). We therefore conducted a Sholl analysis and quantified the number of dendritic intersections at concentric 10 μm intervals from the cell body through the molecular layer. Our initial analysis included all cells that had at least 40% of the dendritic tree length in the analyzed section, and suggested that older-adult-born neurons had more intersections at distal dendritic regions. However, these results were biased by the larger number of cut dendrites in the 16w-neonatal group relative to the adult-born neuron groups. We therefore excluded cells where < 70% of the total dendritic length was present in the analyzed section, and found 10-30 neurons per group that fit this criterion. Younger cells still tended to have more complete neurons but group differences were not statistically significant (% of neuron present in analyzed section: F_4,88_=2.6, P=0.04; post-hoc comparisons all P>0.05; number of cut dendrites: F_4,88_=3.1, P=0.02; post-hoc comparisons all P>0.05). The Sholl analysis revealed that the number of dendritic branch intersections increased at progressively greater distances from the cell soma (Fig. 2d). Neonatal-born neurons had more intersections than adult-born groups at proximal dendritic regions, reflecting their positioning in the superficial granule cell layer, closer to the inner molecular layer. Immature, 2w cells had fewer dendritic intersections in proximal and distal regions, but were not different from the other groups in the intermediate dendritic tree. Proximal and distal dendritic intersections continued to increase from 4-7 weeks of cell age, at which point adult-born neurons were comparable to neonatally-born neurons, aside from having fewer intersections at the proximal dendritic tree.

To obtain a complementary measure of dendritic structure that is independent of dendritic length and is not influenced by tissue sectioning, we quantified the total number of dendritic branches across the full dendritic tree (Fig. 2e). Consistent with recent in vivo and in vitro results (Gonçalves et al., 2016; Beining et al., 2017; Jungenitz et al., 2018), we observed significantly greater numbers of dendritic branches in immature cells. 2w cells had the most dendritic branches; there was significant pruning of branches from 2-4w, and no changes from 4-7w. However, the total number of dendritic branches then increased between 7w and 24w, indicating a later wave of dendritic growth in adult-born neurons.

To identify where dendritic branching differed, we quantified branching according to order, where primary branches are those that emanate directly from the cell body, and the order increases by 1 with each branch point (Fig. 2f). Dendrites typically bifurcated at branch points but trifurcation was also observed in all groups except the 24w group. Overall, 13% of cells trifurcated (2w: 16/77 cells; 4w: 6/49 cells; 7w: 7/64 cells, 24w: 0/37 cells; 16w-neonatal: 14/73 cells; examples in Movie 1). Consistent with the Sholl data, the number of lower-order (1°-3°) branches increased as adult-born cells aged and, by 24w, was comparable to neonatally-born cells. Adult-born neurons are commonly recognized to have a single primary dendrite (Wang et al., 2000; Kerloch et al., 2018), which we observed in 2-7w cells. However, by 24w adult-born cells had, on average, 2 primary dendrites. Quaternary branches were the most common, and did not differ across groups. Whereas the number of lower order dendrites correlated with cell age, the number of higher order dendrites tended to show the opposite pattern. 2w cells had significantly more high-order branches (5°, 6°) compared to all other groups. Neonatal-born neurons had the fewest 5° branches.

Finally, we calculated the branching index of cells, a measure that normalizes dendritic branching to the number of primary dendrites (branch tips / # primary dendrites; Fig. 2g). The branching index was greater in 2w cells than all other groups. In older cells, the distribution of branching indices tended to be bimodal; cells with a single primary dendrite had a branching index that was ∼twice that of cells with ≥ 2 primary dendrites. When we excluded cells that had more than one primary dendrite, 2w cells still had a greater branching index than all older-aged adult-born cells, indicating that their greater branching index is due to more extensive distal branching and not simply because they tend to only have 1 primary dendrite.

Our branch order analyses raise the question of how adult-born neurons gain additional primary dendrites. A second primary dendrite could be generated de novo from the cell body or it could arise by branching off of the existing dendritic tree. It seemed unlikely that additional primary dendrites arise via sprouting because, out of 294 cells examined, only 4 cells possessed an unbranched primary dendrite and only 1 of these cells was adult-born (24w). We therefore hypothesized that new primary dendrites may emerge from the existing dendritic tree, via an “unzipping” of the primary dendrite until the first branch point meets the soma. Indeed, amongst cells with a single primary dendrite, we found a significant shortening between 2 and 4 weeks (Fig. 3a). There were no further changes, possibly because the cells with the shortest primary dendrites become excluded from the analysis as they mature and gain a second primary dendrite.

From 2-24 weeks there was a gradual transition from cells having only a single primary dendrite to having 2 or more primary dendrites (Fig. 3b). Since the biggest transition occurred between 7-24 weeks, we focussed on 7 weeks for further analyses. We reasoned that, if primary dendrites “unzip”, there should be similar patterns of branching in cells with 1 vs 2 primary dendrites. Indeed, when we excluded the primary dendrite from cells with only 1 primary dendrite (1P), the branch order pattern became identical to cells that had 2 primary dendrites (2P; Fig. 3c-d). We then examined the branch index and total length of 1º-branch-trees (i.e. trees associated with a primary dendrite; a 1P cell will have one, and a 2P cell will have two, 1º-branch-trees). For both measures, 1º-branch-trees of 1P cells were double that of 2P cells, consistent with a model where unzipping a longer, more complex 1º-branch-tree results in 2 simpler 1º-branch-trees (Fig. 3e). However, 1º-branch-trees had a much greater range of total lengths and were more variable in 2P cells, suggesting possible within-cell variation in 1º-branch-tree complexity (branch index coefficients of variation (CV): 2P cells: 45%, 1P cells: 14%; length CVs: 2P cells: 34%, 1P cells: 18%). If 2P cells have one 1º-branch-tree that is less developed than the other this could either be due to immaturity (supporting a sprouting model) or it could be an innate property that existed prior to unzipping. If innate, then differences in dendritic complexity should be apparent in the 2º-branch-trees of 1P cells that have not (yet) unzipped. To test this, we compared the branching order, branch index and total length of 1º-branch-trees (2P cells) and 2º-branch-trees (1P cells; Fig, 3f-i). For all measures, there was within-cell variation where a “high” branch-tree had significantly greater complexity and length compared to a “low” branch-tree. This pattern was observed in both 1P cells and 2P cells, supporting a model where inherent differences in dendritic sub-trees exist before a primary dendrite unzips to form 2 primary dendrites in an adult-born neuron (Fig. 3j). Further, the average and minimum total lengths of “low” 1º-branch-trees on 2P cells were 1012 μm and 499 μm, respectively. To sprout trees of these lengths from 4-7w, dendrites would have to grow at rates of 48 and 15 μm/day, respectively, which is unlikely given that the average rate of dendritic growth for the entire dendritic tree over this interval was only 15 μm/day.

### Spines

To assess putative postsynaptic targets of cortical and subcortical axons, protrusion densities on DG neurons were quantified throughout the molecular layer (Fig. 4). The total number of cells examined were: 85 (2w), 46 (4w), 66 (24w), 76 (16w-neonatal). There were few protrusions on 2w cells (0.2/μm) but a dramatic 500% increase between 2-4w and a further 80% increase from 4-7w. Total protrusion densities did not increase further between 7w and 24w, but plateaued at levels that were ∼60% greater than neonatally-born neurons (Fig. 4b).

Protrusion densities for cells of a given age were similar whether calculated by averaging animal means or by pooling and averaging all cells (< 4% difference). The same pattern of differences were also observed when groups were comprised of animal averaged data rather than individual cell data (F_4,54_=205, all comparisons P < 0.0001, except 4w vs 16w-neonatal P=0.08 and 7w vs 24w P=0.4).

Distinct inputs are segregated along granule neuron dendritic trees, where the lateral entorhinal cortex targets the outer molecular layer, the medial entorhinal cortex targets the middle molecular later, and subcortical and commissural fibers target the inner molecular layer (Leranth and Hajszan, 2007; Witter, 2007). We therefore examined whether the maturational profile of postsynaptic sites differs along these functionally-relevant anatomical subregions and found fewer protrusions in the inner molecular layer at 4 and 7 weeks of age (Fig. 4c). This regional difference was absent by 24 weeks of age and not present in neonatally-born neurons, suggesting delayed maturation of commissural and/or subcortical inputs onto adult-born neurons. Consistent with these data, protrusion density was generally not different across dendritic branch orders, except for 2w and 4w cells, which had lower densities on primary and secondary branches compared to their higher order branches (Fig. 4d).

Spines can be categorized into functionally-relevant subclasses based on morphology, where thin filopodial protrusions tend to be transient, plastic potential synaptic partners and large mushroom spines are structurally stable, synaptically stronger, and believed to be sites of long-term information storage (Holtmaat and Svoboda, 2009; Berry and Nedivi, 2017). Consistent with a developmental role, filopodia density peaked at 7 weeks (Fig. 4e). The density of thin spines followed the same pattern as the protrusion densities, and accounted for most of the protrusions (Fig. 4f). We additionally quantified large mushroom spines in different-aged DG neurons (Fig. 4g). Whereas mushroom spines were virtually absent from young 2-week-old cells, they steadily increased with age. At 24 weeks, densities of mushroom spines were greater than all other groups and they were nearly twice as common as in neonatally-born neurons. The regional distribution of mushroom spines resembled the overall spine density pattern, where fewer mushroom spines were found in the inner molecular layer on 4- and 7-week-old neurons (Fig. 4h). Here, however, there were also fewer mushroom spines observed on neonatally-born neurons in the inner molecular layer. Examining the changes in proportion of spine type revealed a clear maturational profile, where filopodia made up the largest proportion of protrusions at 2 weeks and declined with age; mushroom spines followed the opposite pattern (Fig. 4i).

### Presynaptic terminals

Dentate granule neuron MFBs are large excitatory presynaptic structures, composed of multiple active zones that target CA3 pyramidal neurons and mossy cells (Chicurel and Harris, 1992; Rollenhagen et al., 2007). Associated with each MFB are smaller filopodial extensions that form synapses with inhibitory neurons (Acsády et al., 1998). To gain insights into the role that cell age may play in efferent connectivity, we first quantified the maximal 2D area of MFBs as an anatomical proxy for synaptic strength. The total number of MFBs examined were: 143 (2w), 95 (4w), 170 (7w), 105 (24w), 200 (16w-neonatal). Among adult-born neurons, MFBs doubled in size between 2-7w, with most growth occurring between 4-7w. However, they grew an additional 20% between 7w and 24w. At 24w, adult-born MFBs were 34% larger than 16w-neonatal-born neurons (Fig. 5b). Across all cell populations, MFBs were smaller in CA3c than in CA3a/b (Fig. 5c).

**Figure 5:**
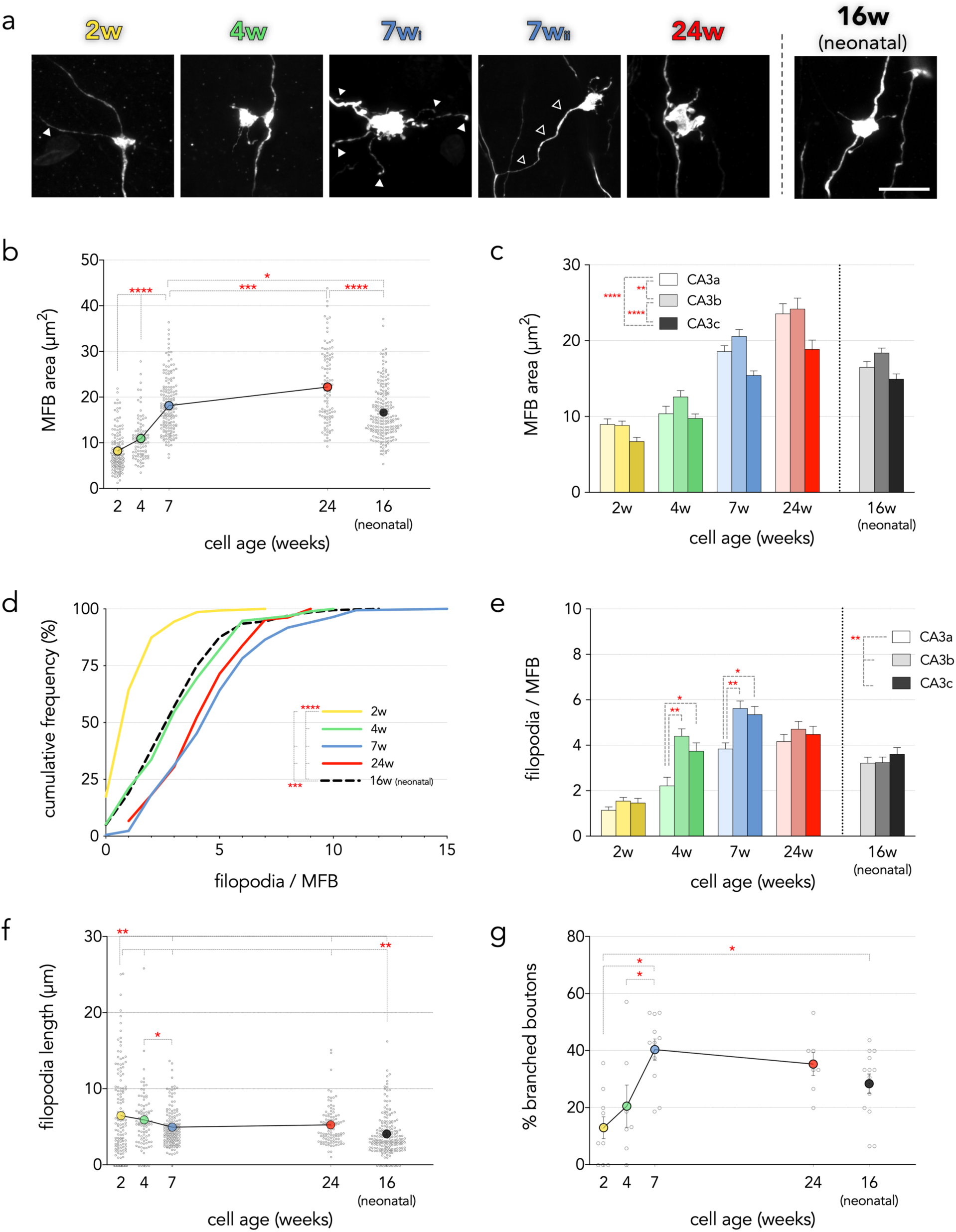
Efferent synaptic terminals of neonatally- and adult-born neurons. a) Confocal images of retrovirally-labelled mossy fiber boutons (MFB) and filopodial terminals. Filled arrowheads in 2w and 7w_i_ images indicate filopodial extensions. Open arrowheads in 7w_ii_ indicate a branched MFB. Scale bar, 10 μm for all images except for 7w_ii_, 11.7 μm. b) MFBs increased in size with cell age (F_4,708_=160, P<0.0001). c) MFBs increased in size from CA3c < CA3a < CA3b (effect of subregion F_2,698_=26, P<0.0001; cell group x subregion interaction F_8,698_=1.2, P=0.3). d) The number of filopodia per MFB increased from 2-7w and remained greater than neonatally-born neurons at 24w (Kruskal-Wallis test, P<0.0001). e) The number of filopodia per MFB was lowest in CA3a, specifically in 4w and 7w cells (effect of subregion F_2,698_=15, P<0.0001; cell group x subregion interaction F_8,698_=2.8, P=0.005). f) Filopodia length was greatest at 2w and decreased with cell age but remained longer than neonatally-born neurons (F_4,667_=18, P<0.0001). g) The proportion of branched MFBs increased with cell age and did not differ significantly between older adult-born neurons and neonatal-born neurons (F_5,47_=6, P<0.001). *P<0.05, **P<0.01, ***P<0.001, ****P<0.0001.

MFB sizes that were calculated from animal averages differed by ≤ 5% from means that were calculated by pooling all cells within a given age group. The same pattern of group differences were also observed when animal averaged data were analyzed rather than individual cell data (F_4,43_=45, all groups significantly different from one another (P<0.05) except 7w vs 16w-neonatal (P=0.13).

We next examined the filopodial processes that protrude off of MFBs and contact GABAergic interneurons. There was an age-related increase in filopodia/MFB from 1.4 filopodia/MFB at 2 weeks to 4-5 filopodia/MFB at 7/24 weeks, which was significantly greater than that of neonatally-born cells (3.4; Fig. 5d). There were also fewer filopodia/MFB in CA3a, an effect that was driven by 4w and 7w cells (Fig. 5e). While young 2-week-old cells had few filopodia/MFB, their filopodia were significantly longer than those of older adult-born and neonatally-born neurons (Fig. 5f). Filopodial length declined from a mean of 7 μm at 2 weeks to ∼5 μm at 7 weeks of age, which was not different from 24-week-old adult-born neurons. Filopodia length of neonatally-born neurons was shorter than all adult-born populations. Filopodia were longer in CA3c than in CA3a, an effect that was not specific to any subpopulation of cells (effect of subregion F_2,657_=4.8, P=0.008; subregion x cell age interaction, F_8,657_=1.4, P=0.2).

In addition to age-related changes in MFB size, there were also differences in the positioning of MFBs relative to the axon (Fig. 5g). Young adult-born cells, 2-4 weeks old, tended to have en passant MFBs that were located directly on the axon. In contrast, 30-40% of MFBs on older adult-born cells and neonatally-born cells were connected to the axon by a branch. Branch lengths averaged 8 μm across all cells examined. 75% of branches were less than 10 μm in length; the longest branch was 52 μm. There were no differences in branch length across cell groups or CA3 subregions (group effect F_4,207_=1.8, P=0.12; CA3 subregion effect F_2,207_=0.1, P=0.9; interaction F_8,207_=0.9, P=0.5).

### Soma and nuclear size

Our analyses indicate an extended developmental trajectory for adult-born neurons, where several morphological features ultimately surpass neonatally-born neurons in size. We tested the generality of our findings by measuring the size of the cell soma and nucleus, both of which vary as adult-born neurons mature (Kirn et al., 1991; van Praag et al., 2002; Amrein and Slomianka, 2010; Radic et al., 2015; Moreno-Jiménez et al., 2019)(Fig. 6). Consistent with early growth, the cell soma increased in size from 2-7 weeks and was stable thereafter and similar to neonatal-born neurons. In contrast, the nuclear size was generally smaller in adult-born cells than neonatal-born cells, except at 7 weeks when adult-born cells had larger nuclei that matched 16w-neonatal cells in size.

**Figure 6:**
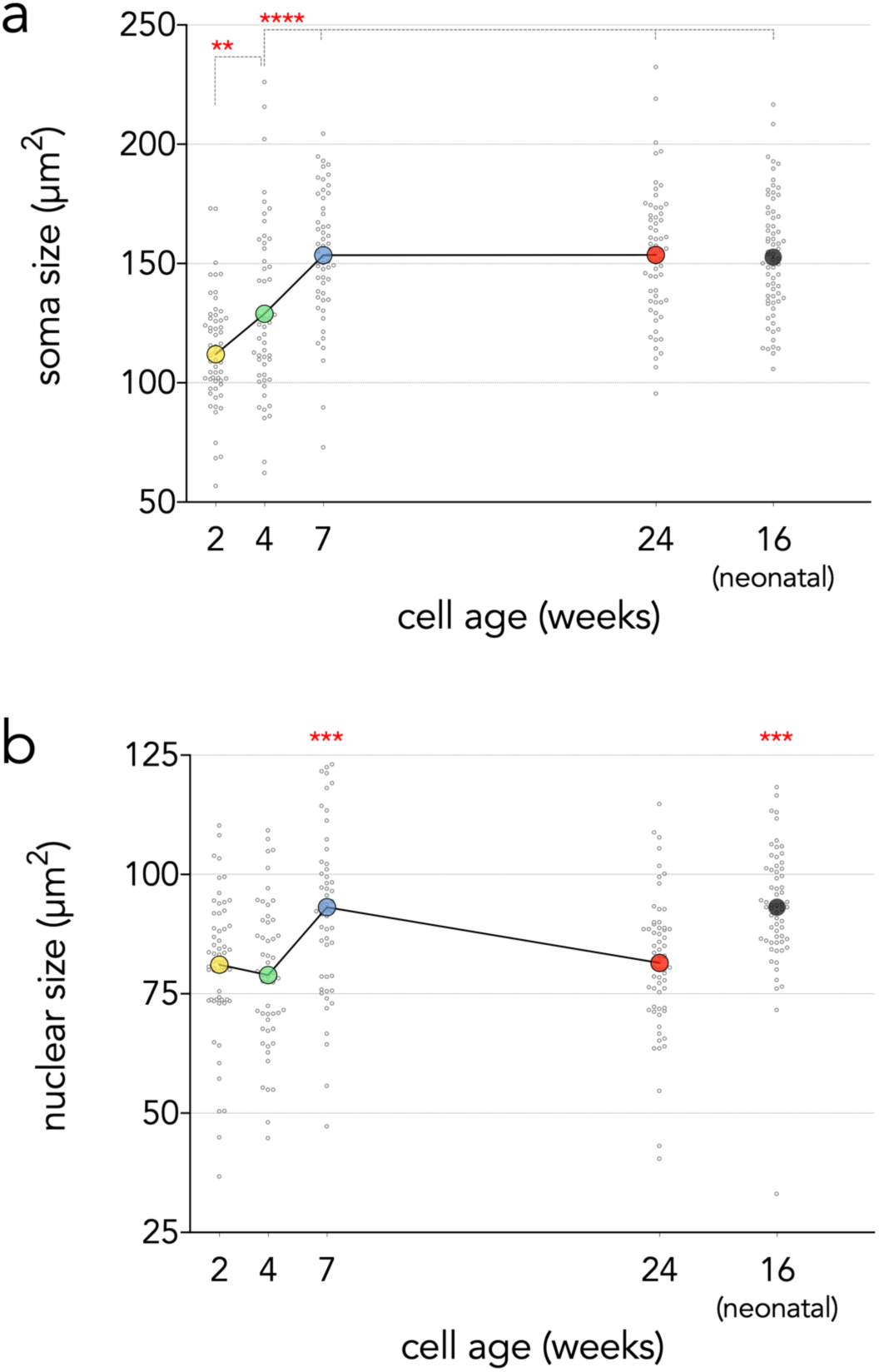
Soma and nuclear morphology. a) The cell soma increased in size as adult-born cells aged, plateauing and matching neonatal-born neurons by 7 weeks (F_4,289_=27, P<0.0001; **P<0.01, ****P<0.0001). b) The nuclear size of adult-born neurons was generally consistent across cell ages, except at 7w when nuclear were larger and equivalent to neonatal-born neurons (F_4,291_=12.5, P<0.0001). ***P<0.001 vs 2w, 4w and 24w.

### Morphological differences are not due to animal age

The morphology of developmentally-born granule cells differs depending on whether cells were born embryonically or neonatally (Kerloch et al., 2018), and differences in experience-dependent gene expression have been observed in cells born at different stages of adulthood (Tronel et al., 2015; Ohline et al., 2018). Thus, it is possible that the observed morphological differences are due to the age of the animal at the time of cell birth, and not the age of the cells. To test this we injected groups of rats with retrovirus at 8 weeks or 14 weeks of age (since this bookends the full range of ages used for our analyses of adult-born cells) and examined cells 7 weeks later (Fig. 7). We observed no differences between 8w-born and 14w-born cells for any of our key measures (dendritic length and branching, spine densities, MFB morphology). We did find that branched MFBs had longer branches in 14w-born cells than in 8w-born cells, suggesting that terminals from later-born neurons may search a greater space to find their postsynaptic targets (Mann-Whitney U=298, *P=0.03). These data collectively indicate that morphological differences between our adult-born cell populations reflect differences in cell age rather than animal age.

**Figure 7:**
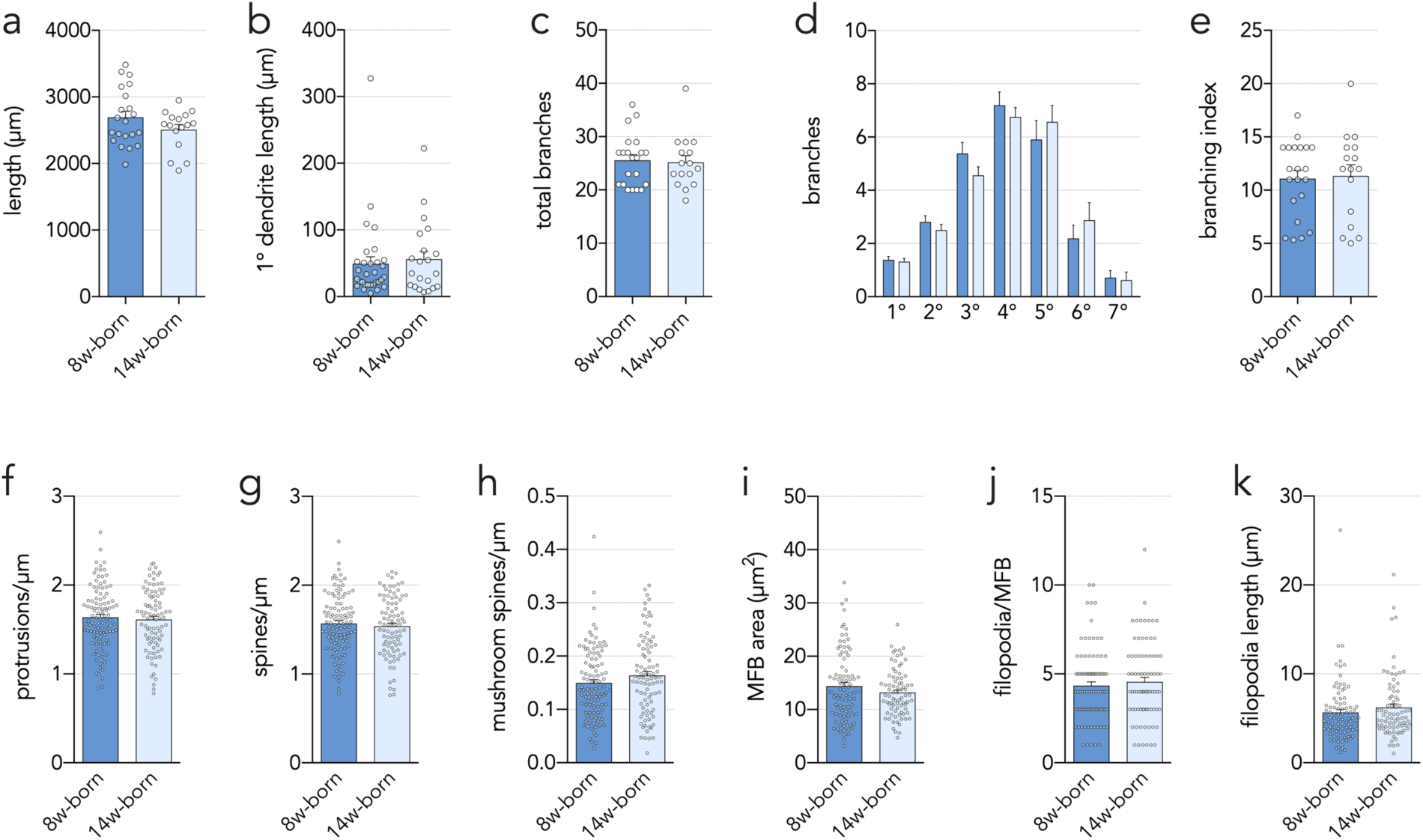
Equivalent morphology of neurons born at different stages of young adulthood. a-e) Morphology did not differ between 7-week-old neurons that were born in 8- vs 14-week-old rats. a) Total dendritic length (T_35_=1.5, P=0.15). b) Primary dendrite length (T_28_=0.5, P=0.63). c) Total number of dendritic branches (T_21_=0.9, P=0.4). d) Branch order distribution (effect of branch order: F_6,126_=35, P<0.0001; effect of birthdate: F_1,21_=0.8, P=0.4; interaction: F_6,126_=0.5, P=0.8). e) Branching index (T_21_=0.3, P=0.8). f) Dendritic protrusions (T_193_=0.5, P=0.6). g) Spines (T_193_=0.7, P=0.5). h) Mushroom spines (T_193_=1.4, P=0.2). i) MFB area (T_167_=0.6, P=0.6). j) Number of MFB filopodia (Mann-Whitney U=3311, P=0.5). k) Filopodia length (T_166_=1.2, P=0.2).

### Modelling the cumulative effects of neuron addition and extended maturation

Given the dramatic decline in neurogenesis with age, it has remained unclear whether new neurons contribute to the function of the aging brain. Our results suggest that extended growth may contribute to the plasticity of the aging brain. Whereas our analyses were limited to individual cells born at a snapshot in time, an accurate representation of plasticity must account for the extended growth of all cells born throughout adult life. To estimate this we created a model that combined and integrated rates of cell addition, dendritic growth and spine growth from 8 weeks to 2 years of age in the rat (see Methods; functions in Fig. 8a-d). Consistent with previous estimates based on different datasets (Snyder and Cameron, 2012), and evidence from Glast CreERT2 mice (DeCarolis et al., 2013), our model predicts that ∼50% of total DG neurons are added in adulthood (Fig. 8e). A similar proportion of dendritic material in the DG can also be attributed to adult-born neurons (∼4km in absolute terms). Given that adult-born neurons achieve greater spine densities than neonatally-born neurons, we estimate that adult neurogenesis ultimately contributes the majority of spines in the DG (Fig. 8g-h). Notably, while neurogenesis in our model ended by 1.5 years, adult-born neurons continued to add to the total dendrite length and total number of spines in the DG even after this point; this was particularly salient for mushroom spines. Between 1-2 years we estimate that adult neurogenesis adds only 2.5% of total cells, consistent with previous quantitative estimates of neurogenesis rates later in life (Lazic, 2012), but during this time it contributes 7% of total dendritic length, 10% of total spines and 24% of mature mushroom spines in the DG.

**Figure 8:**
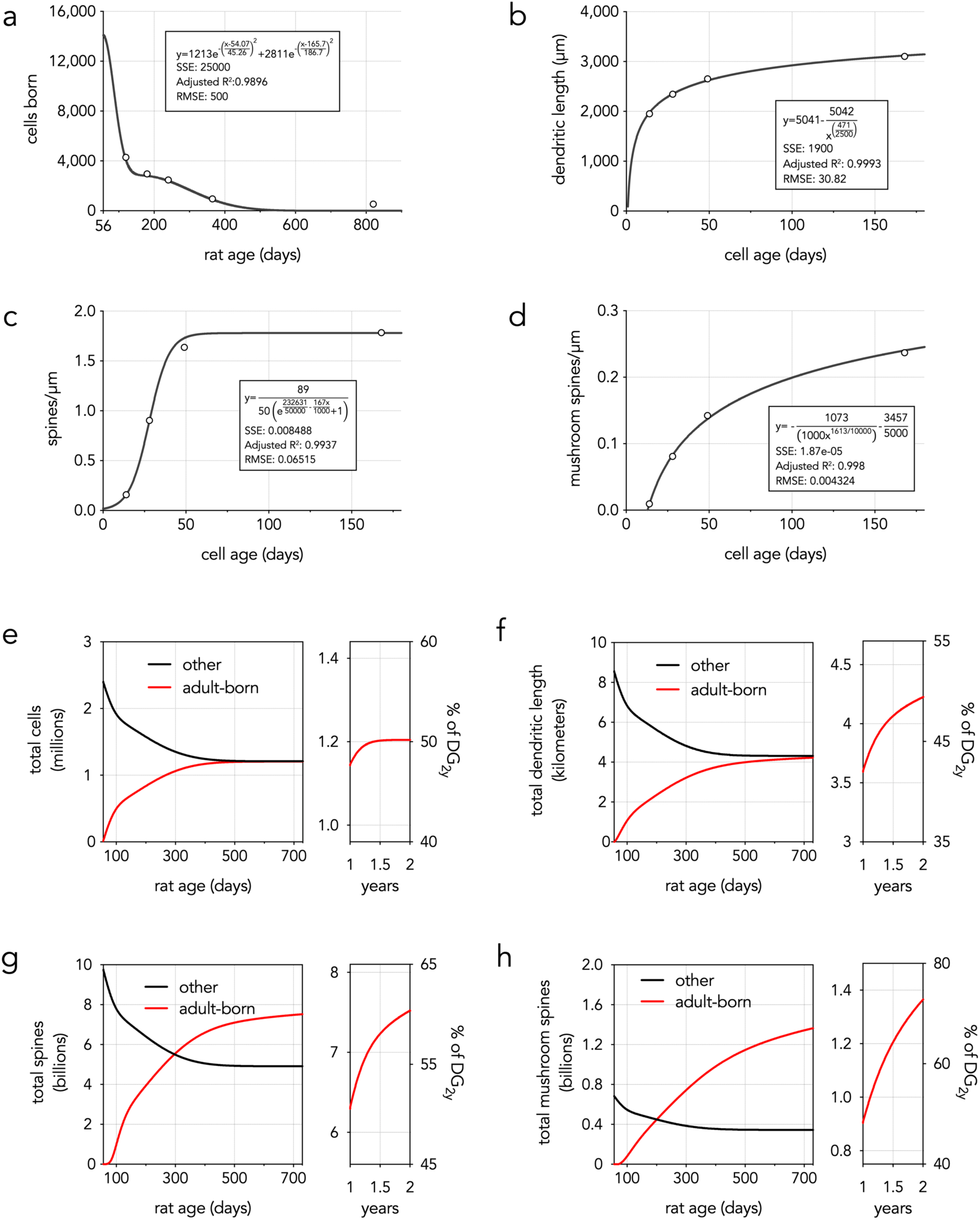
Modelling cumulative effects of adult neurogenesis. a-d) Functions used to model neuron addition in adulthood (a), dendritic growth of adult-born neurons (b), spine growth on adult-born neurons (c) and mushroom spine growth on adult-born neurons (d). e-h) Modelling neurogenic plasticity throughout life. Square left panels show neurogenesis effects across the full period of adulthood of a rat (8 weeks to 2 years). Narrow right panels focus on cumulative effects during the 2^nd^ half of life, when neurogenesis has declined but growth continues; right y-axis shows percent contribution by neurogenesis, relative to the total granule cell population at 2 years of life. Other/adult-born = cells born before/after 8 weeks of age. e) Adult-born neurons accumulate throughout the first 1.5 years of life and ultimately account for ∼half the total granule cell population (numbers are bilateral. f) Total dendritic length contributed by adult neurogenesis increases throughout life, even though neurogenesis ends at 1.5 years in this model. Due to prolonged growth of dendrites and spines, the total number of spines (thin + mushroom; g) and mushroom spines (h) added to the dentate gyrus increases through old age.

## DISCUSSION

Here we report that functionally-relevant morphological features of adult-born neurons develop over a surprisingly long timeframe. From 2-7 weeks, neurons displayed classic patterns of morphological maturation. However, properties of 7w cells suggest a later wave of development: dendrites were thickest, nuclei were largest, and dendritic filopodia were present in greatest numbers. By examining cells at 24 weeks, which is substantially older than in most previous studies, we found that dendrites, spines and presynaptic terminals continued to grow and even surpass those of neonatally-born neurons. Our results are not confounded by ontogeny since neurons of the same age, born in 8- and 14-week-old rats, had equivalent morphology. Differences between 24w adult-born neurons and 16w neonatal-born neurons are not simply due to differences in cell age (where 24w adult-born neurons had more time to grow) because adult-born cells already had greater spine densities, larger MFBs, and more MFB filopodia by 7w. In fact, developmentally-born cells mature faster than adult-born neurons (Overstreet-Wadiche et al., 2006; Zhao et al., 2006; Cahill et al., 2017) and do not undergo additional dendritic growth beyond ∼5-6 weeks of cell age (Kerloch et al., 2018). 16w-neonatal neurons therefore reflect a relevant control population of mature neurons, though it will be important for future studies to characterize the maturational trajectory of dendrites, spines and terminals on developmentally-born neurons. While physiological experiments are needed to determine functional significance, our results indicate that adult-born neurons are plastic well beyond the traditional critical window and may make unique contributions to hippocampal functioning for the lifespan of the cell.

### Extended growth of adult-born neuron dendrites

Immature neurons displayed a transient overproduction of distal, high-order branches as has been previously described in vivo (Gonçalves et al., 2016) and in vitro (Beining et al., 2017). While branches were pruned between 2-4 weeks, total dendritic length grew significantly during this interval and dendrites continued to grow until 24w. Why hasn’t the continued growth of adult-born neuronal dendrites (beyond ∼2 months) been reported previously? The most likely explanation is that most studies have not investigated older cohorts of cells (Shapiro et al., 2007; Wang et al., 2008; Leslie et al., 2011; Piatti et al., 2011; Dieni et al., 2013). While some studies have analyzed cells as old as 8-11w, small sample sizes or cut dendrites may have masked growth (Sun et al., 2013; Gonçalves et al., 2016; Beining et al., 2017; Trinchero et al., 2017; Bolós et al., 2019). Quantifying growth over an extended interval of 17w may also have facilitated detection of cumulative growth that is not readily apparent over shorter intervals. Whether growth is linear between 7-24w, and whether adult-born cells continue to grow beyond 24w, remain open questions.

Whereas prolonged dendritic growth suggests a long-term plasticity function for adult-born neurons, their branching pattern matures to become broadly similar to neonatally-born neurons. Typically, adult-born neurons are considered to have only a single primary dendrite (Wang et al., 2000; Kerloch et al., 2018). However, forced swimming, inflammation, GSK-3 overexpression and traumatic brain injury can increase the number of primary dendrites, indicating an unusual form of dendritic plasticity (Llorens-Martín et al., 2014; Villasana et al., 2015; Llorens-Martín et al., 2016). By examining branching patterns at early and late ages, our data suggests an “unzipping” model where the first branch point of the primary dendrite moves closer to the soma with age (or the soma moves towards the branch point). Subtrees with variable length and branching complexity suggest that the granule cell dendritic tree is computationally compartmentalized (Losonczy et al., 2008). Furthermore, at least some mature neonatal- and adult-born neurons have morphological features of highly-active granule neurons (few primary dendrites and extensive higher-order branching (Diamantaki et al., 2016)).

### Prolonged maturation and high density of dendritic spines

The gradual loss of immature filopodia and acquisition of mature mushroom spines is consistent with previous evidence that spine morphology changes with adult-born cell age (Toni et al., 2007). While the inner molecular layer is the site of the first synapses onto adult-born neurons (Chancey et al., 2014), our data suggest that synaptic maturation in this region is delayed at 7 weeks but may be accelerated by a single day of water maze training. Consistent with our data, previous reports have found that adult-born neurons gain spines beyond 1 month (van Praag et al., 2002; Zhao et al., 2006; Jessberger et al., 2007; Toni et al., 2007; Jungenitz et al., 2018; Bolós et al., 2019) and display experience-dependent changes in spines and connectivity when cells are several months old (Lemaire et al., 2012; Bergami et al., 2015). We found that adult-born neurons ultimately achieved greater spine density than neonatally-born neurons, which has likely gone unnoticed since, to our knowledge, only one other study has used retrovirus to compare spine densities of developmentally- and adult-born neurons (Toni et al., 2007). While that study also reported an extended period of spine formation and maturation, spine densities on old adult-born neurons (180 days) and developmentally-born (P4) neurons were similar. The reason for this discrepancy is unclear but may stem from any number of methodological differences (e.g. species, sex, running, cell birthdate). Regardless, the higher spine density of adult-born neurons is consistent with Golgi studies of mice, primates and humans that identified a subpopulation of DG granule neurons that have ∼2x the normal density of dendritic spines (Williams and Matthysse, 1983; Seress and Frotscher, 1990; Seress, 1992).

### Protracted growth of large mossy fiber boutons

Mossy fiber boutons are large multisynapse complexes that target the proximal dendrites of CA3 pyramidal neurons (Amaral and Dent, 1981; Chicurel and Harris, 1992; Rollenhagen et al., 2007) and are theorized to play a dominant role in recruiting CA3 pyramidal neurons during memory encoding (Rolls, 2010). Indeed, a single mossy fiber can trigger spiking in postsynaptic pyramidal neurons (Henze et al., 2002; Vyleta et al., 2016). Retroviral studies have found that MFB size, the number of active zones and the number of synaptic vesicles increases as adult-born cells mature over 2-10 weeks of age (Faulkner et al., 2008; Toni et al., 2008; Restivo et al., 2015; Bolós et al., 2019). Given the positive relationship between MFB and EPSP size (Galimberti et al., 2006; 2010), our data suggests that adult neurogenesis contributes to the heterogeneity of synaptic strength at the DG-CA3 synapse, and may produce a population of particularly powerful synapses. Since adult-born neurons initially share CA3 spines with existing neurons before developing fully independent synapses (Toni et al., 2008), and neonatally-born neurons turn over in significant numbers (Dayer et al., 2003; Cahill et al., 2017; Ciric et al., 2019), adult-born neurons may outcompete developmentally-born neurons for CA3 circuit connectivity.

A portion of MFBs were not directly attached to the main axon but instead were connected via a small branch. Similar “terminal boutons” in the neocortex are more morphologically plastic than en passant boutons (De Paola et al., 2006), suggesting that branched boutons may play a unique role in hippocampal function. Branching is likely to influence signal propagation and coding properties of axons (Ofer et al., 2017) and, since voltage-gated channels are differentially distributed across axonal compartments, branches could also offer an anatomical substrate for modulating the active properties of mossy fibers (Engel and Jonas, 2005; Kole et al., 2008; Rowan et al., 2016).

Thin filopodial protrusions extend off of MFBs to excite inhibitory interneurons (Acsády et al., 1998; McBain, 2008) and adult-born neurons play an important role in recruiting inhibitory networks (Drew et al., 2015; Restivo et al., 2015). Consistent with these data, we found that the number of filopodia per MFB plateaued at 7 weeks and remained greater than developmentally-born neurons at 24 weeks. Thus, adult-born neurons may play a long-term role in shaping inhibition in CA3, which could promote memory precision (Ruediger et al., 2011; Guo et al., 2018) by reducing overlap between ensembles of neurons that represent different experiences (Niibori et al., 2012).

### Significance of extended development over the lifespan

Critical periods endow adult-born neurons with a unique capacity for plasticity during their immature stages. However, the focus on immature neurons has come at the expense of understanding the properties of older neurons, and led to the assumption that adult-born neurons lose their functional relevance with age (Snyder, 2019). By integrating the extended growth of adult-born neurons into a model of neuronal accumulation, we present evidence that neurogenesis makes a dramatic contribution to the overall structural plasticity of the dentate gyrus. Our model predicts the addition of 1.2 billion spines (10% of the total) between 1-2 years, and 300 million spines (2.5% of the total) between 1.5-2 years, which is after cell proliferation has ended. Since human granule neurons grow throughout middle to old age (Flood et al., 1985; Coleman and Flood, 1987), and neonatally-born neurons in mice do not grow over similar intervals (Kerloch et al., 2018), adult-born neurons may offer a unique reserve of plasticity in aging (Kempermann, 2008), when the medial-temporal lobe becomes vulnerable to pathology (Leal and Yassa, 2015). Moreover, cellular maturation is slower in older (Trinchero et al., 2017) and longer lived animals (Kohler et al., 2011), which may further prolong neurogenesis-associated plasticity in human aging. It will be important for future work to further characterize this long-term plasticity in cells from female subjects and in other DG subregions (e.g. infrapyramidal blade, ventral DG).

Plasticity aside, our data are consistent with emerging evidence that adult-born granule neurons are functionally distinct from neurons born at other stages of life, even after they have “matured” (Snyder, 2019). The significantly greater spine density suggests that older adult-born neurons may be inherently more likely to associate, and be recruited by, cortical inputs. The presence of larger boutons suggests that adult-born neurons may be more capable of depolarizing postsynaptic pyramidal neurons, and the presence of more filopodia suggests they may be more effective at refining CA3 representations through feedforward inhibition. Given cellular heterogeneity in disease vulnerability, protracted neurogenesis may also result in subpopulations of cells that are differentially susceptible to pathology.

## Acknowledgements

This work was supported by the Natural Sciences and Engineering Research Council of Canada, the Canadian Institutes of Health Research and the Michael Smith Foundation for Health Research.

## MOVIE LEGEND

<u>Movie 1: dendrite trifurcation</u>. This video shows confocal z-stack animations that focus through dendritic trees of 2w, 4w, 7w and 16w-neonatal cells. Arrows indicate points where dendrites trifurcate.

## EXTENDED DATA LEGEND

Figure 2-1: Raw data. This spreadsheet contains the raw data.

